# Enhanced recognition of topoisomerase 1 upon DNA binding by a subset of anti-topoisomerase 1 autoantibodies in systemic sclerosis

**DOI:** 10.1101/2025.05.26.656088

**Authors:** Sam Neppelenbroek, Corrie M. Wortel, Danique M.H. van Rijswijck, Marjolein J.A.L. Wetzels, Robbert Q. Kim, Nivine E.W. Levarht, Albert J.R. Heck, Jeska K. de Vries-Bouwstra, Cynthia M. Fehres, René E.M. Toes, Hans U. Scherer

**Affiliations:** Department of Rheumatology, Leiden University Medical Center, Albinusdreef 2, 2333 ZA, Leiden, The Netherlands; Biomolecular Mass Spectrometry and Proteomics, Bijvoet Center for Biomolecular Research and Utrecht Institute for Pharmaceutical Sciences, Utrecht University, Padualaan 8, 3584 CH, Utrecht, The Netherlands; Protein Facility, Department of Cell and Chemical Biology, Leiden University Medical Center, Einthovenweg 20, 2333 ZC, Leiden, The Netherlands

## Abstract

**Objectives:** Systemic sclerosis (SSc) is a deleterious disease. Its clinical management is complicated by strong interpatient heterogeneity. Progressive organ fibrosis is linked to autoantibodies against topoisomerase 1 (TOP1). Here, we hypothesized that the molecular interaction between anti-TOP1 autoantibodies (ATAs) and TOP1 could be a relevant determinant of SSc pathogenesis. DNA binding by TOP1 might affect its interaction with ATAs. Therefore, we studied the effect of DNA binding by TOP1 on ATA recognition.

**Methods:** ATA monoclonal antibodies (ATA mAbs) were generated from patient-derived TOP1-reactive B cells. Reactivity of ATA mAbs and antibodies in patient plasma towards TOP1 and TOP1-DNA cleavage complexes (TOP1cc) was determined by ELISA and mass photometry. Immunostimulatory properties of ATA mAbs in complex with TOP1/TOP1cc were assessed by stimulation of monocytic THP-1 cells.

**Results:** DNA binding by TOP1 differentially affected the recognition of TOP1 by ATA mAbs. A subset of mAbs showed enhanced binding to TOP1cc, whereas others recognized TOP1 and TOP1cc to a similar extent. ATAs with enhanced TOP1cc recognition were observed in plasma of ATA^+^ SSc patients and correlated with ATA levels and interstitial lung disease. These ‘TOP1cc-enhanced ATAs’ variably affected the enzymatic function of TOP1 and circulated predominantly as IgG1 and IgM. The interaction of ‘TOP1cc-enhanced ATA mAbs’ with TOP1cc strongly increased IL-8 production by THP-1 cells.

**Conclusions:** The differential recognition of TOP1 upon DNA binding by ATAs demonstrates heterogeneity in the ATA B cell response, possibly impacting on disease-relevant processes in severe SSc.

## Introduction

Systemic sclerosis (SSc) is a heterogeneous disease characterized by fibrosis of internal organs and the skin, vasculopathy and autoimmunity. The disease displays remarkable clinical heterogeneity ranging from indolent to rapidly progressive disease with irreversible organ damage and subsequent mortality. Clinical management of SSc is complicated by the lack of effective treatments and variation in disease progression, which is difficult to predict^1^. Therefore, identification of pathogenic mechanisms driving disease may improve patient stratification and enable the development of targeted treatment strategies.

Like many other autoimmune diseases, SSc is hallmarked by a break in B cell tolerance towards nuclear proteins^2,3^. Autoantibodies targeting the nuclear enzyme topoisomerase 1 (TOP1) are observed in a subset of SSc patients and associate with diffuse cutaneous SSc, a distinct clinical phenotype with the highest mortality among the rheumatic diseases^4,5^. Specifically, anti-topoisomerase 1 autoantibodies (ATAs) associate with progressive lung and skin fibrosis and early death, although not all ATA-positive patients show this progressive course of disease^6^. Recently, we could associate measures reflective of an active ATA B cell response to disease progression in ATA^+^ SSc patients^7,8^. Specifically, ATA-secreting B cells were found in the circulation of SSc patients and their presence associated with the presence and severity of pulmonary fibrosis^7^. Also, the presence of ATA-IgM was predictive of disease progression in the early stages of disease^8^. These findings suggest that activated, autoreactive ATA B cells might contribute directly to processes driving fibrosis. This notion is further supported by the beneficial effects of B cell-targeting therapies in SSc^9,10^. Nevertheless, the mechanisms by which ATA B cells could contribute to disease remain unclear^3^. ATA pathogenicity has been debated, with the hypothesis that ATAs directly activate fibroblasts^11–13^. However, whether this affects disease *in vivo* is unclear, primarily because ATA can also be present in indolent disease. ATA B cells, on the other hand, might contribute to disease by the production of cytokines and stimulation of T cells through antigen presentation and co-stimulation^14^.

A crucial prerequisite for B cell activation is the recognition of antigen by B cell receptors (BCR)^15,16^. The same holds for antibody-mediated effector functions which require antibody-binding to cognate antigens. Epitope structure and conformation can determine and modulate these interactions^17^. TOP1 is a DNA-binding nuclear protein which unwinds negatively supercoiled DNA^18^. During its enzymatic cycle, TOP1 covalently binds DNA to form a TOP1-DNA cleavage complex (TOP1cc) which allows the passing of the other DNA strand through the single-stranded break^18^. TOP1cc is a transient complex under physiological conditions, but stable TOP1cc can be formed if DNA contains lesions, which can occur during cell death induced under inflammatory conditions^19^. Binding of antigen to nucleic acids, a common property among many nuclear antigens targeted in autoimmune diseases, can potentiate antibody effector functions and B cell activation through co-engagement of innate Toll-like receptors with Fc receptors or the BCR^2,16^. Whether TOP1 as SSc autoantigen binds DNA, and how TOP1 and TOP1cc are recognized by human ATAs, warrants detailed analyses to better understand the antigenicity of TOP1 in SSc pathogenesis.

To this end, we set out to determine the molecular and functional characteristics of the interaction between ATAs and TOP1. We generated patient-derived monoclonal ATAs (ATA mAbs) enabling us to study the effect of DNA binding by TOP1 on the interaction with ATAs. We found remarkable heterogeneity in the recognition of TOP1 and TOP1cc among ATA mAbs and ATA in plasma. This heterogeneity affected antibody functionality and likely impacts on disease-relevant processes in progressive SSc. Together, these findings contribute to our understanding of disease variability in ATA^+^ SSc.

## Methods

The experimental procedures for the recombinant production and purification of human TOP1, the generation of patient-derived ATA mAbs, gel electrophoresis, western blotting, cross-inhibition ELISA with ATA mAbs, mass photometry and TOP1-mediated DNA relaxation assay are provided in detail in the supplementary methods section.

### Patients and healthy individuals

Peripheral blood was obtained from ATA^+^ and anti-centromere protein B autoantibody^+^ (ACA^+^) SSc patients visiting the outpatient clinic of the Department of Rheumatology at Leiden University Medical Center. These SSc patients were included in the Leiden Combined Care in Systemic Sclerosis (CCISS) cohort^20^, fulfilled the ACR/EULAR 2013 classification criteria for SSc and were either ATA-IgG^+^ or ACA-IgG^+^ as measured by routine clinical diagnostic tests. Patients on B-cell depleting therapies or patients with a history of hematopoietic stem cell transplantation were excluded. Plasma samples of a cross-sectional cohort of ATA-IgG^+^ SSc patients (n=21) collected in a previous study were used (Table 1)^7^. In addition, serum samples of ATA-IgG^+^ SSc patients containing high levels of ATA-IgM (n=9) were selected from another previously reported study (Table 1)^8^. Plasma samples from ACA^+^ SSc patients (n=9) and healthy donors (n=6) were used as controls (Table 1).

**Table 1.** Characteristics of ATA^+^ SSc patients.

|  | ATA-IgG<sup>+</sup> SSc patients<br>(n=21) | ATA-IgM<sup>high</sup> SSc patients<br>(n=9) | ACA-IgG<sup>+</sup> SSc patients<br>(n=9) | Healthy donors<br>(n=6) |
| --- | --- | --- | --- | --- |
| Age, years, median (IQR) | 55 (39-65) | 53 (36-67) | 55 (46-68) | 29 (26-36) |
| Sex, female, n (%) | 11 (52%) | 8 (89%) | 9 (100%) | 5 (83%) |
| Duration since first NRP symptom, months, median (IQR) | 56 (31-92.8) | 48 (14.5-119) | 117 (57-359) |  |
| Diffuse cutaneous SSc, n (%) | 7 (33%) | 3 (33%) | 0 (0%) |  |
| mRSS, median (IQR) | 5 (2-7.5) | 6 (4-16) | 5 (0-9) |  |
| ILD on HRCT, n (%) | 14 (66.7%) | 8 (89%) | 0 (0%) |  |
| Current use of immunosuppressive drugs, n (%) | 12 (57%) | 1 (11%) | 3 (33%) |  |

IQR = interquartile range, NRP = non-Raynaud's phenomenon, mRSS = modified Rodnan skin score, ILD = interstitial lung disease, HRCT = high-resolution computer tomography scan.

### Generation of TOP1cc using suicide substrate

To obtain stable TOP1-DNA cleavage complexes (TOP1cc), TOP1 was incubated with a suicide substrate (ScS) which is covalently bound by TOP1. Suicide substrate was generated as described previously by Kristoffersen *et al*^21^. Three oligonucleotides (26DL: ‘5-pAGA AAA ATT TTT CCG AGT GCG AAG-3’ (p = phosphorylated 5’), 38SC: 5’-CTA GAG GAT CTA AAA GAC TTA GA-3’, 61NCL: 5’-CTT CGC ACT CGG AAA AAT TTT TCT AAG TCT TTT AGA TCC TCT AG-3’) were mixed to obtain a 66.67µM concentration of ScS. The suspension was heated to 80°C for 5 minutes and cooled to room temperature in 3 hours. TOP1cc was formed by incubation of 0.1mg/ml of TOP1 with 5µM of ScS for 30 minutes at 37°C.

### Anti-TOP1/TOP1cc-antibody ELISAs

To measure the reactivity of ATA mAbs and antibodies in plasma towards TOP1 and TOP1cc, high binding 384-well microplates (Corning) were coated with 2.5µg/ml TOP1 or TOP1cc in PBS (pH=8). Plates were blocked using 1% BSA in PBS (Ig, IgG, IgG subclasses and IgA) or 1% skim milk in PBS (IgM). Plasma, ATA mAbs and culture supernatants of single cell cultured B cells were applied in single or serial dilutions (as indicated in the respective figures) in 1% BSA in PBS with 0.05% Tween (Ig, IgG, IgG subclasses and IgA) or in 1% skim milk in PBS with 0.05% Tween (IgM). Subsequently, secondary antibodies were added: rabbit anti-human IgG-HRP (1:5000, P0214, DAKO), mouse anti-human IgG1 (1:5000, HP6186, Nordic-MUbio), mouse anti-human IgG2 (1:250, HP6014, Nordic-MUbio), mouse anti-human IgG3 (HP6080, 1:2500, Nordic-MUbio), mouse anti-human IgG4 (1:500, MH164-1, Sanquin), goat anti-human IgA-HRP (1:1000, A18781, Novex), goat anti-human-IgM-HRP (1:5000, AP114P, Millipore), and goat anti-human Ig-HRP (1:10000, A80-152P, Bethyl Laboratories). Goat anti-mouse Ig-HRP (1:5000 for IgG1/2, 1:2500 for IgG3/4, P0447, DAKO) was used to detect the IgG subclass-specific, secondary antibodies. Pooled plasma samples were used as reference standards. ATA mAbs were used in all plasma ELISAs to control for comparable coating of TOP1 and TOP1cc (ATA mAb 2F8) and as positive controls for the differential recognition of TOP1cc compared to TOP1 (ATA mAbs 7G6 and 9D11). ELISAs were developed using ABTS supplemented with H_2_O_2_. Optical density was measured at 415 nanometers with a Spectramax I3x Multi-Mode Microplate Reader (Molecular Devices) using SoftMax® Pro 7 (version 7.0.2, Molecular Devices).

### Stimulation of THP-1 cells

To assess the immunostimulatory properties of ATA mAbs, monocytic THP-1 cells were stimulated with ATA mAbs in complex with plate-bound TOP1 or TOP1cc. NUNC Maxisorp 96-well plates (ThermoFisher Scientific) were coated with 5µg/ml TOP1 and TOP1cc. Plates were blocked with PBS 1% BSA and 2.5µg/ml of LPS-depleted (Pierce^TM^ High Capacity Endotoxin Removal Spin Columns, ThermoFisher Scientific), sterile-filtered ATA mAbs were subsequently added to the plates. The anti-citrullinated protein mAb 1F2 was used as negative control^22^. Upon the formation of plate-bound immune complexes, 1×10^5^ THP1 cells were added to the plates and stimulated for 24 hours at 37°C, 5% CO2. Stimulation with 1µg/ml lipopolysaccharide (LPS, Sigma) served as positive control. Upon stimulation, culture supernatants were collected to determine the secretion of IL-8 by the THP-1 cells.

### IL-8 ELISA

Human IL-8 was measured using a commercial ELISA on culture supernatants of THP-1 cells in accordance with the manufacturer’s instructions (DY208-05, R&D Systems). Volumes of the ELISA were adapted for the use of high binding 384- well Microplates (Corning): 15µl for coating, sample, and BD OptEIA^TM^ TMB substrate solution (555214, BD Biosciences), 75µl for blocking (1% BSA in PBS), and 7.5µl for stop solution (1M H_2_SO_4_). Reactions were developed for 50 minutes and absorbance was measured with a Spectramax I3x Multi-Mode Microplate Reader (Molecular Devices) at 450 nanometers with correction of the absorbance at 570nm.

### Statistical analysis

Data were analyzed using Graphpad Prism (version 10.2.3). As indicated in the figure legends, Mann-Whitney U test, Wilcoxon matched-pairs signed rank test, and Kruskal-Wallis test combined with Dunn’s multiple comparison were used where appropriate to determine the statistical significance of the data. Correlations were described using Spearman’s rank correlation coefficient.

### Study approval

This study was approved by the ethical review board of the Leiden University Medical Center (protocol number P17.151) and the Leiden University Biobank Toetsing Commissie (REU/036/SH/sh). Included patients gave written informed consent to participate in the study.

## Results

### Generation and characterization of patient-derived anti-topoisomerase 1 monoclonal antibodies

We first generated ATA mAbs based on BCR sequences of TOP1-reactive B cells obtained from an ATA^+^ SSc patient. B cells from peripheral blood of the patient were single cell sorted based on their reactivity towards TOP1 conjugated to Alexa Fluor^TM^ 647 and phycoerythrin (Fig. 1A). These sorted cells were subsequently differentiated towards antibody-secreting cells in culture. Secreted antibodies could be detected in a fraction of the culture supernatants (Suppl. Fig. 1A-C). By analyzing the reactivity of the secreted antibodies towards human TOP1, we identified several TOP1-reactive clones (Fig. 1B-D). BCR sequences were determined from three of these clones (termed 2F8, 7G6 and 9D11; Table 2) and expressed recombinantly as IgG1 mAbs (Suppl. Fig. 1D). Reactivity of the generated ATA mAbs towards TOP1 was verified by ELISA in a concentration-dependent manner (Fig. 1E). TOP1 binding by ATA-IgG 2F8 was stronger compared to ATA-IgGs 7G6 and 9D11. Specificity of ATA mAbs was further confirmed by western blotting, with all ATA mAbs recognizing one band at the expected height of TOP1 (Fig. 1F, Suppl. Fig. 1E). No binding was observed to centromeric protein B (CENP-B) and double-stranded DNA (Fig. 1G-H).

**Figure 1.**
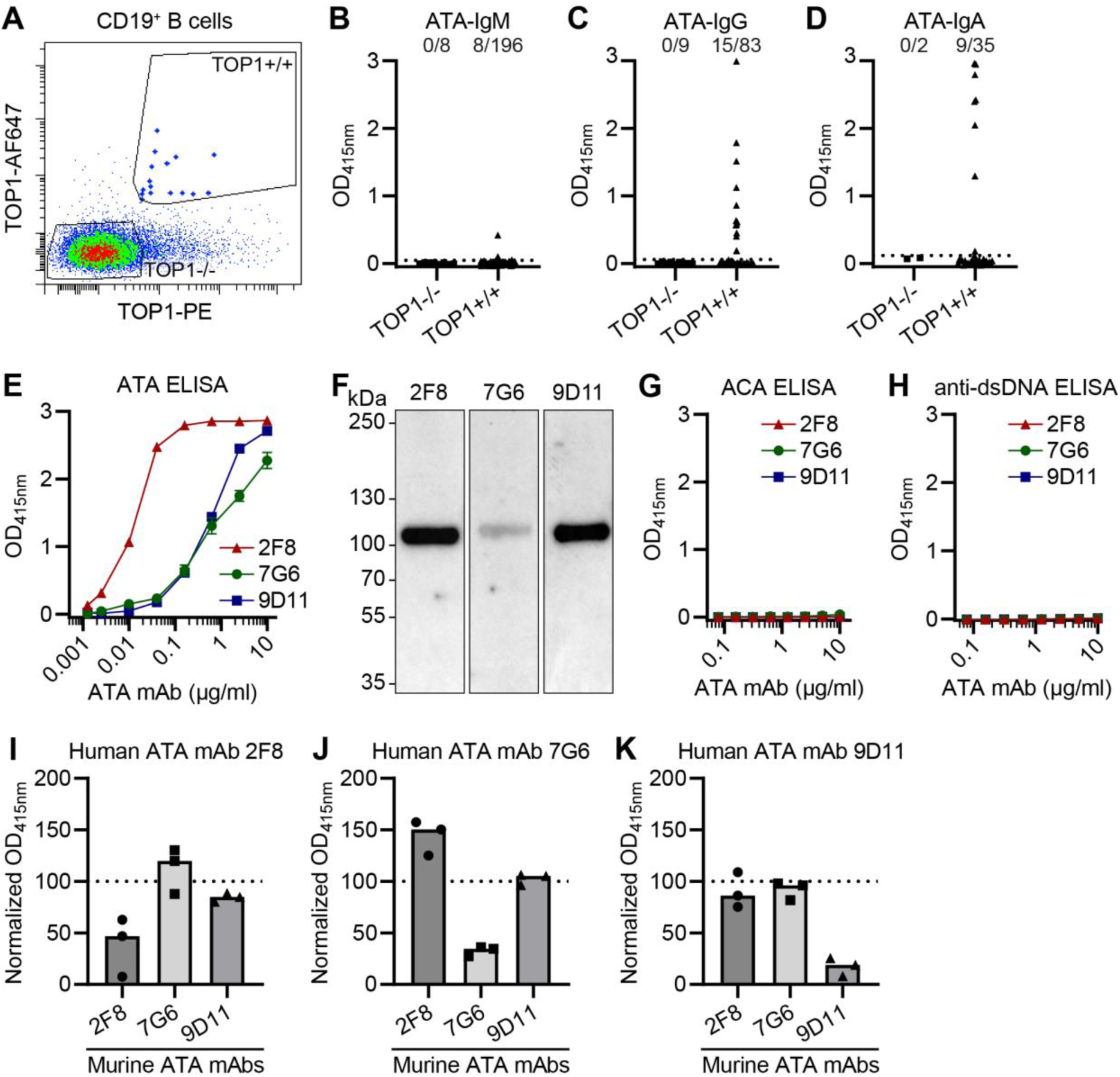
Generation and characterization of patient-derived ATA mAbs. **(A)** Gating strategy used to single-cell sort CD19^+^ TOP1^+/+^ B cells binding to TOP1-Alexa Fluor^TM^ 647 (AF647) and TOP1-phycoerythrin (PE). **(B-D)** Reactivity of IgM (B), IgG (C) and IgA (D) secreted by single-cell-sorted TOP1^+/+^ B cells towards TOP1 as assessed by an anti-TOP1-Ig (ATA-IgM), anti-TOP1-IgG and anti-TOP1-IgA ELISA, respectively. Cut-off (mean TOP1^-/-^ B cells + 4x standard deviation) is represented by dotted line. Only culture supernatants in which IgM, IgG or IgA could be detected (Suppl. Fig. 1A-C) are depicted. **(E)** Reactivity of patient-derived ATA mAbs towards TOP1 in an ATA-IgG ELISA. **(F)** Binding of TOP1 by ATA mAbs on western blot. Molecular weight in kilodaltons (kDa) was estimated using a protein ladder. **(G-H)** Reactivity of ATA mAbs towards centromere protein B in an anti-centromere protein B antibody-IgG (ACA-IgG) ELISA (G) and towards double-stranded DNA in an anti-double-stranded DNA-IgG (anti-dsDNA-IgG) ELISA (H). **(I-K)** Binding of ATA mAbs 2F8 (I), 7G6 (J) and 9D11 (K) with a human constant domain (human ATA mAbs) to a TOP1-coated ELISA plate pre-incubated with 5µg/ml of the indicated ATA mAbs with a murine constant domain (murine ATA mAbs). Binding was normalized to condition in which no murine mAb was added. Concentrations of human mAbs were in the linear range of the ATA-IgG ELISA; 2F8 - 0.04µg/ml, 7G6 & 9D11 - 1µg/ml. Data is derived from three independent experiments. (B-E, G-K) Optical density was measured at 415 nanometers (OD_415nm_). (E, G-H) Error bars represent standard deviation of technical duplicates. (E-H) Data are representative of at least two independent experiments.

**Table 2.**
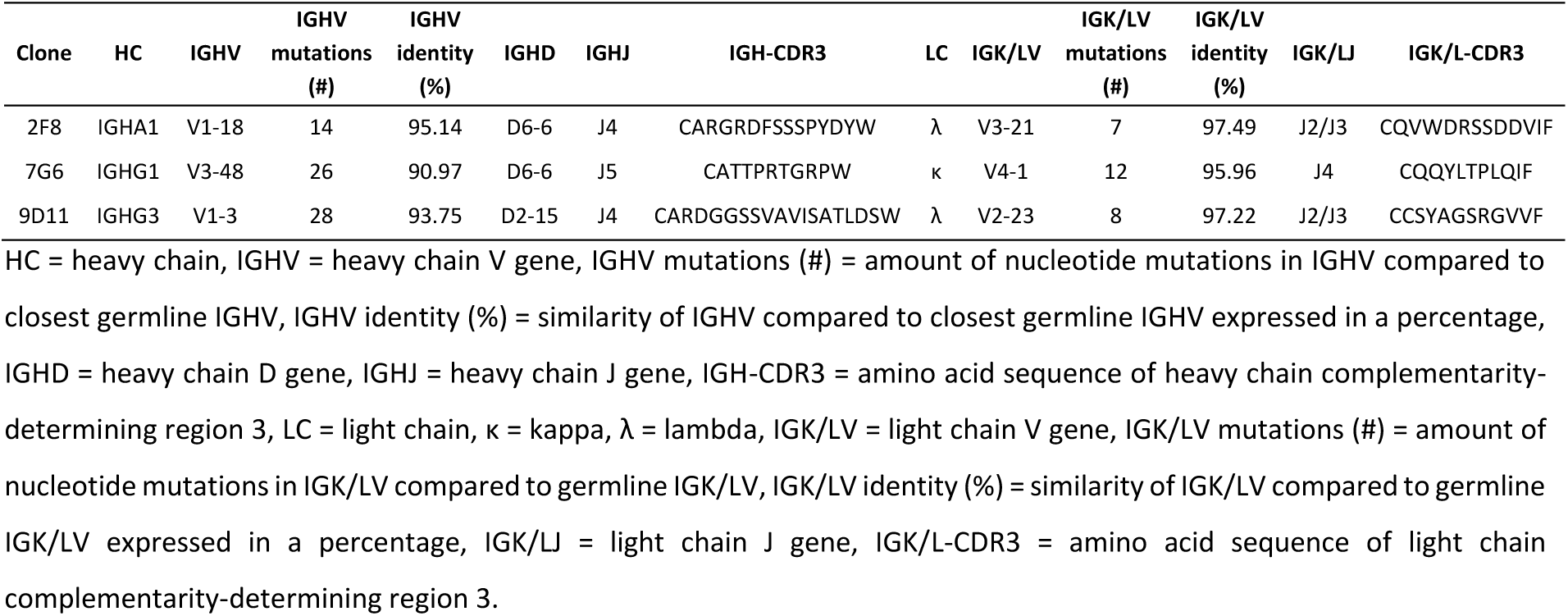
B cell receptor sequence details of patient-derived ATA mAbs.

We also performed cross-inhibition assays to determine whether the recombinantly produced ATA mAbs would bind unique or overlapping TOP1 epitopes. The three ATA mAbs were generated with murine constant domains to test their capacity to inhibit TOP1 binding by ATA mAbs with a human constant domain. Binding of human ATA mAbs to TOP1 could be blocked by their direct murine counterparts, but not by the respective other mAbs (Fig. 1I-K, Suppl. Fig. 1F). Together, these data show that we generated three ATA mAbs binding unique epitopes of human TOP1.

### A subset of ATA mAbs recognize TOP1 better in complex with DNA

TOP1 can form a covalent complex with DNA, termed TOP1-DNA cleavage complex (TOP1cc)^18^. To study the effect of DNA binding on the recognition of TOP1 by the ATA mAbs, we next compared mAb reactivity towards TOP1 and TOP1cc. TOP1cc was formed using a suicide substrate, a double-stranded DNA ligand which covalently and irreversibly binds to TOP1^21^. This binding results in a size shift which can be visualized by gel electrophoresis (Fig. 2A). By coating of either antigen on an ELISA plate, the recognition of TOP1 and TOP1cc by the ATA mAbs could be compared. ATA mAb 2F8 recognized TOP1 and TOP1cc to a similar extent (Fig. 2B-C). In contrast, ATA mAbs 7G6 and 9D11 recognized TOP1cc with higher relative avidity than TOP1 (Fig. 2B-C). None of the mAbs reacted to suicide substrate alone (Suppl. Fig. 2A). To further substantiate the differential binding of ATA to TOP1 and TOP1cc, we measured the reactivity towards either antigen by IgG in culture supernatants of 13 additional, single-cell sorted TOP1-reactive B cell clones. Only 3 of the 13 clones recognized TOP1 and TOP1cc to a similar extent, whereas the other 10 clones showed enhanced recognition of TOP1cc as compared to TOP1 (Fig. 2D). Importantly, we noted a strong variation between clones as to the extent to which TOP1cc was differentially recognized.

**Figure 2.**
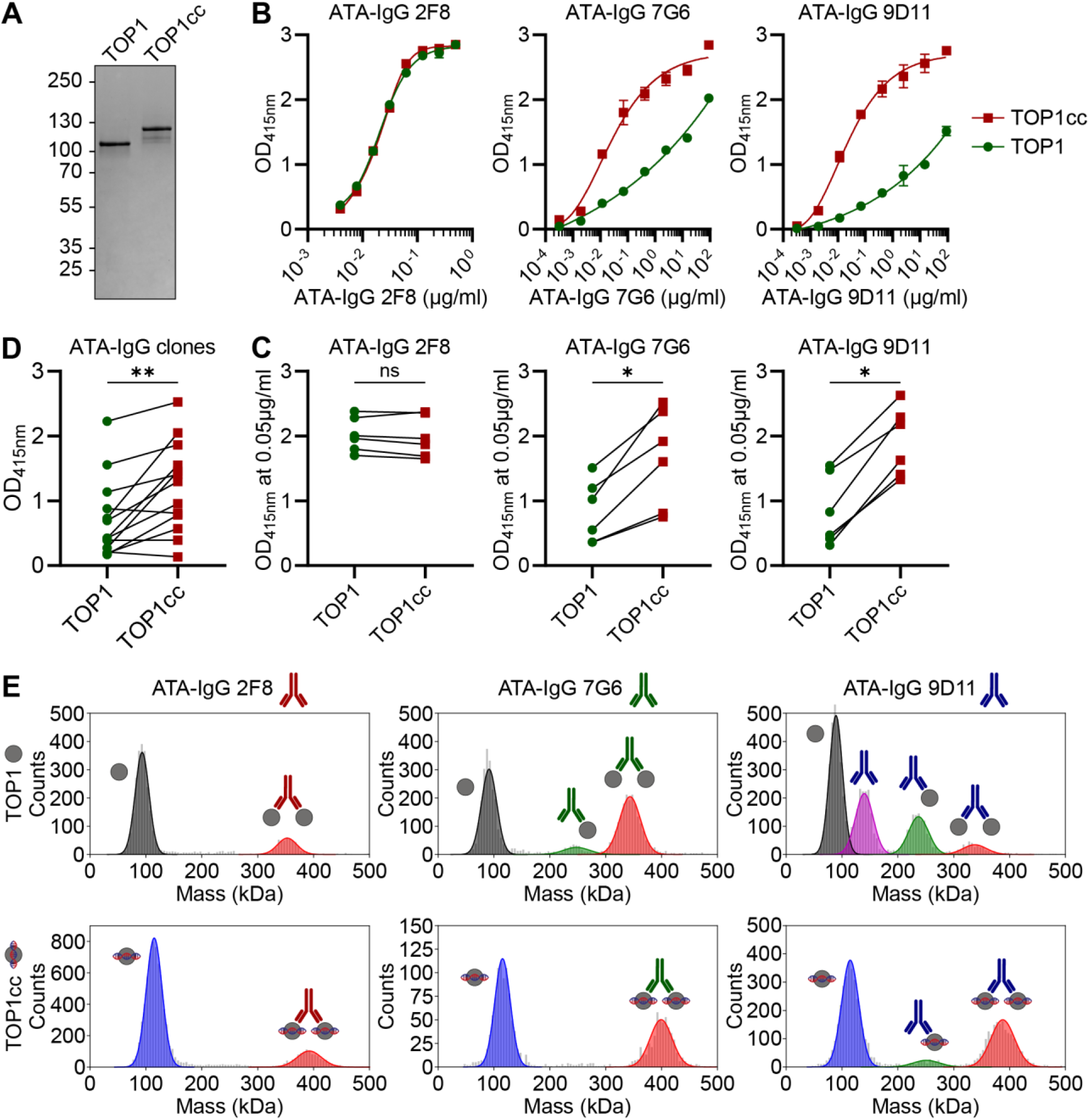
Reactivity of ATA mAbs towards TOP1 and TOP1cc. **(A)** Gel electrophoresis of TOP1 upon pre-incubation without or with suicide substrate (TOP1 and TOP1cc, respectively) under non-reducing conditions. Molecular weight in kilodaltons (kDa) was estimated using a protein ladder. **(B)** Reactivity of ATA mAbs 2F8, 7G6 and 9D11 towards TOP1 and TOP1cc in an anti-TOP1/TOP1cc-IgG ELISA. Data points represent the mean and standard deviation of two technical replicates from an individual experiment. **(C)** Quantification of the binding of 0.05µg/ml ATA mAbs 2F8, 7G6 and 9D11 to TOP1 and TOP1cc. Data is derived from six independent experiments. **(D)** Reactivity towards TOP1 and TOP1cc by IgG in cultures supernatants (n=13) of single-cell sorted CD19^+^ TOP1^+/+^ B cells with verified reactivity towards TOP1. **(E)** Representative mass histograms of ATA mAbs incubated with an excess of TOP1 and TOP1cc as obtained by mass photometry. Data is representative of two independent experiments. An overview of detected masses can be found in Suppl. Table 1. (B-D) Optical density was measured at 415 nanometers (OD_415nm_). (C-D) Wilcoxon matched-pairs signed rank test was used to test for significant differences. ns = not significant, * = p<0.05, ** = p<0.005, *** = p<0.001

We next analyzed the differential binding under native conditions in solution to exclude that immobilization of either protein to the ELISA plate would have caused the observed effects. We employed mass photometry to determine the binding stoichiometry of the ATA mAbs to TOP1 and TOP1cc. This single particle interferometric scattering microscopy technique allows to analyze the formation of immune complexes in solution formed upon incubation of ATA mAbs with TOP1 and TOP1cc (Fig. 2E, Suppl. Fig. 2B-C, Suppl. Table 1). In the presence of antigen excess, ATA-IgG 2F8 bound both TOP1 and TOP1cc and efficiently formed immune complexes containing two antigens (Fig. 2E). ATA-IgG 7G6 and ATA-IgG 9D11 did not show complete binding to TOP1 (Fig. 2E). In these conditions, antibodies binding no or only one antigen could be detected, especially for ATA-IgG 9D11. In contrast, nearly complete binding of 7G6 and 9D11 was observed to TOP1cc (Fig. 2E). Hence, this independent approach confirmed the finding that certain ATA mAbs differentially recognize TOP1 and TOP1cc.

### Polyclonal IgG autoantibodies from ATA^+^ SSc patients differentially recognize TOP1cc

Differential recognition of TOP1/TOP1cc by ATA-IgG could affect the functionality of ATAs and, hence, influence disease pathogenesis. Therefore, we next assessed differential recognition of TOP1/TOP1cc by polyclonal ATA in the circulation of ATA^+^ SSc patients. Half-maximal binding titers of polyclonal IgG to TOP1 and TOP1cc were determined by serial dilutions of patient plasma (Fig. 3A). ATA mAbs were taken along in these assays to control for comparable coating concentrations of TOP1 and TOP1cc (ATA mAb 2F8), and as positive controls for the differential recognition of TOP1cc compared to TOP1 (ATA mAbs 7G6 and 9D11). Plasma of all ATA^+^ SSc patients (n=21) showed higher IgG titers to TOP1cc compared to TOP1 (Fig. 3B). Patients varied as to the extent of differential TOP1/TOP1cc recognition. The difference between anti-TOP1-IgG and anti-TOP1cc-IgG titers was modest for some patients. For others, much higher dilutions of plasma were needed to reach the same binding signal for TOP1cc in comparison to TOP1 (up to 30.000 difference in dilution factor) (Fig. 3C). Again, we excluded suicide substrate recognition as confounder, as binding to suicide substrate was minimal to undetectable (Suppl. Fig. 3A). Also, plasma from ACA^+^ SSc patients and healthy donors generated signals below the limit of detection (Suppl. Fig. 3B). Together, these findings indicate that enhanced TOP1cc recognition by a subset of ATAs is a feature commonly present in ATA^+^ SSc patients.

**Figure 3.**
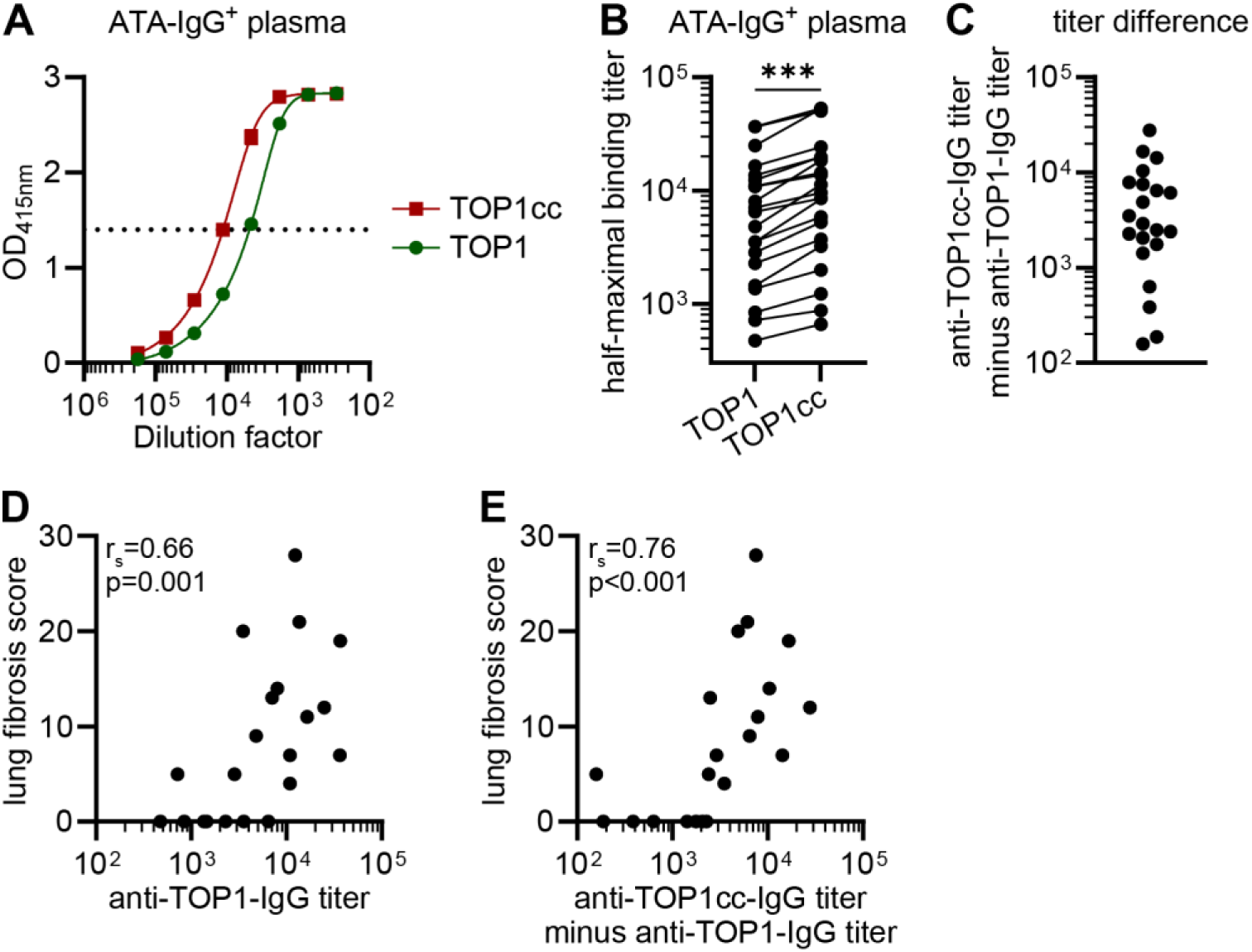
Reactivity of IgG in ATA-IgG^+^ plasma towards TOP1 and TOP1cc. **(A)** Binding of IgG in plasma of an ATA^+^ SSc patient to TOP1 and TOP1cc in an anti-TOP1/TOP1cc-IgG ELISA. Optical density was measured at 415 nanometers (OD_415nm_). Dotted line represents an OD_415nm_ of 1.4 which was considered the half-maximal signal of the assay. **(B)** Anti-TOP1/TOP1cc-IgG expressed as half-maximal binding titers (interpolated dilution generating an optical density of 1.4 at 415 nanometers) in plasma of ATA^+^ SSc patients (n=21). Half-maximal binding titers were statistically compared using the Wilcoxon matched-pairs signed rank test. ns = not significant, * = p<0.05, ** = p<0.005, *** = p<0.001. **(C)** Difference between anti-TOP1cc-IgG titer and anti-TOP1-IgG titer in plasma of ATA^+^ SSc patients. (**D-E**) Correlation between the lung fibrosis score and the anti-TOP1-IgG titer (D) and the difference between anti-TOP1cc-IgG and anti-TOP1-IgG titer (E). Correlations are described using the Spearman’s rank correlation coefficient.

To explore the possible clinical relevance of our finding, we assessed the relation between the differential recognition of TOP1cc and clinical disease parameters using the absolute difference between anti-TOP1cc-IgG and anti-TOP1-IgG titers (anti-TOP1cc-IgG titer minus anti-TOP1-IgG titer) as read-out. This difference correlated with anti-TOP1-IgG titers (r_s_=0.91) (Suppl. Fig. 3C). In addition, both the anti-TOP1-IgG titer and the difference between anti-TOP1cc-IgG and anti-TOP1-IgG titers correlated with the lung fibrosis score (r_s_=0.65 and r_s_=0.76, respectively) (Fig. 3D-E). Similar trends were observed for the modified Rodnan skin score (Suppl. Fig. 3D-E). Together, the data indicate that anti-TOP1-IgG titers and anti-TOP1cc-IgG titers are strongly related and correlate with clinical disease.

### Anti-TOP1cc autoantibodies are predominantly present in the IgG1 and IgM compartment

Isotype and subclass usage strongly affects antibody effector functions and provides insights into the underlying B cell response^23,24^. Therefore, we studied the recognition of TOP1 and TOP1cc by antibodies of various subclasses and isotypes in plasma of ATA^+^ SSc patients. Presence of ATA among the IgG subclasses was variable, as IgG1, IgG2, IgG3 and IgG4 anti-TOP1-autoantibodies could be detected in 67%, 81%, 52% and 95% of the samples (n=21), respectively (Suppl. Fig. 4A-D). Therefore, we limited the analysis to samples in which TOP1 was recognized by multiple IgG subclasses (n=8 for IgG1/3/4, n=7 for IgG2) and in which total IgG recognized TOP1cc better than TOP1 (Fig. 4A). Enhanced recognition of TOP1cc compared to TOP1 was observed for IgG1 and IgG4, but not for IgG2 and IgG3 (Fig. 4A). To compare the relative difference in TOP1cc binding by the IgG subclasses to total IgG, anti-TOP1cc titers were normalized to anti-TOP1 titers by expressing the titers as a ratio (anti-TOP1cc titer divided by anti-TOP1 titer). The relative difference in TOP1cc binding by IgG1 was comparable to total IgG, while the ratio between anti-TOP1cc and anti-TOP1 titers for IgG2-4 were lower (Fig. 4B). Consequently, the differential recognition of TOP1 upon DNA binding appears most prominent for IgG1.

**Figure 4.**
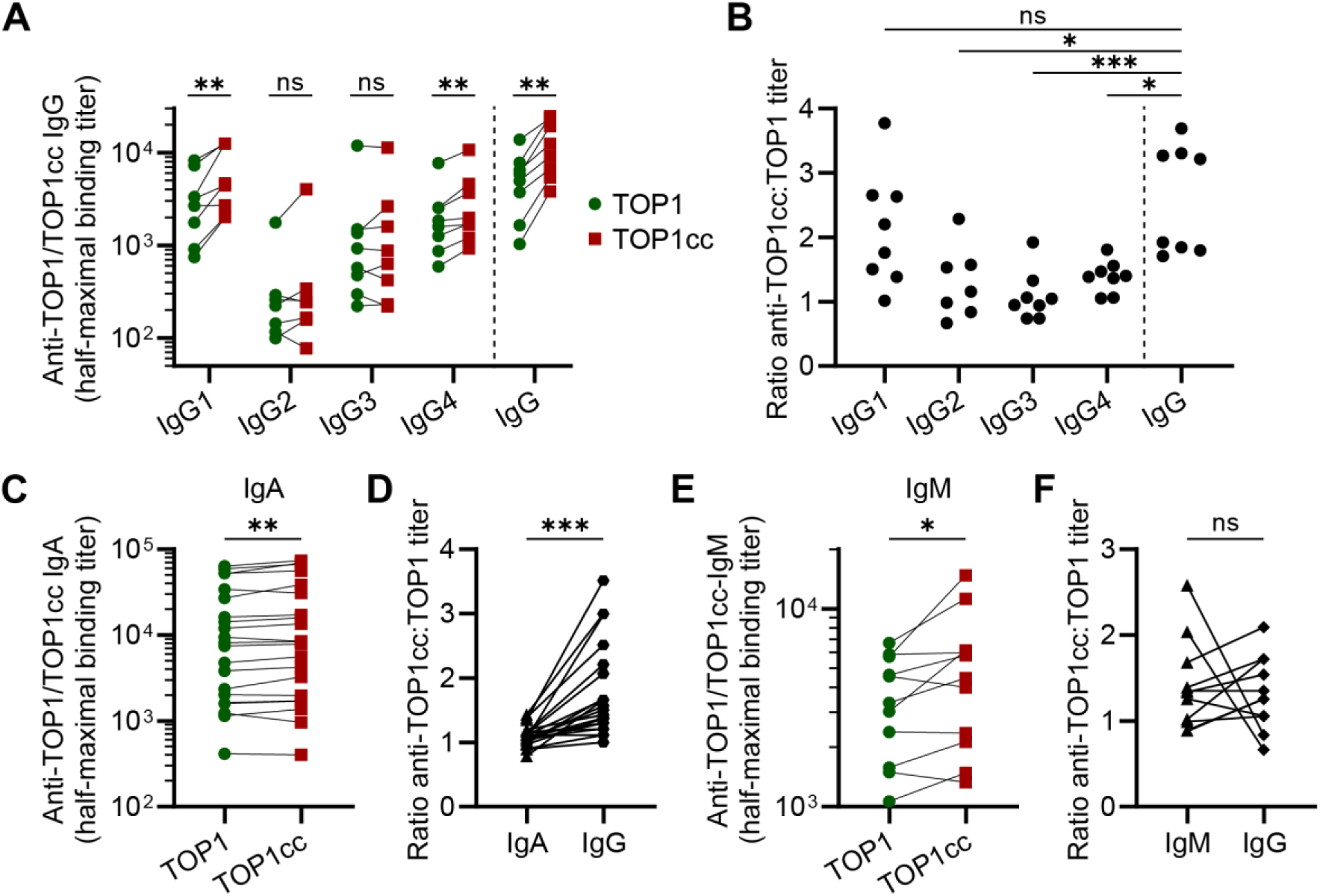
Reactivity towards TOP1 and TOP1cc by IgG subclasses, IgA and IgM in plasma of ATA^+^ SSc patients. **(A)** Half-maximal binding titers of IgG1-4 and total IgG in plasma of ATA-IgG^+^ SSc patients (n=8 for IgG1, 3, 4, n=7 for IgG2) to TOP1 and TOP1cc. **(B)** Ratio between anti-TOP1cc and anti-TOP1 titers of IgG1-4 and total IgG in plasma. Ratios of IgG1-4 were compared to total IgG using the Kruskal-Wallis test combined with Dunn’s multiple comparison. **(C)** Reactivity of IgA in plasma of ATA-IgG^+^ SSc patients (n=21) towards TOP1 and TOP1cc. **(D)** Anti-TOP1cc-IgA/G titers expressed as ratio to anti-TOP1-IgA/G titers. **(E)** Reactivity of IgM in plasma of ATA-IgM^+^ SSc patients (n=11) towards TOP1 and TOP1cc. **(F)** Anti-TOP1cc-IgM/IgG titers expressed as ratio to anti-TOP1-IgA/G titers. (A, C, E) Binding to TOP1 and TOP1cc is expressed as half-maximal binding titer, the interpolated dilution generating an optical density of 1.4 at 415 nanometers. (C-E) Half-maximal binding titers to TOP1 and TOP1cc were statistically compared using the Wilcoxon matched-pairs signed rank test. (D, F) Ratio titers were compared using the Mann-Whitney U test. (A-F) ns = not significant, * = p<0.05, ** = p<0.005, *** = p<0.001.

TOP1-reactive antibodies of the IgA isotype can be detected in almost all ATA-IgG^+^ SSc patients^8^. Accordingly, we detected anti-TOP1-IgA in all ATA-IgG^+^ SSc patients (n=21) included in this study (Fig. 4C). These IgA antibodies recognized TOP1cc slightly better in comparison to TOP1 (Fig. 4C). However, the relative difference in the half-maximal binding titers between TOP1 and TOP1cc was significantly higher in the IgG compartment (Fig. 4D).

For IgM, we observed that only 2 out of 21 ATA-IgG^+^ patients generated higher signals on the anti-TOP1-IgM ELISA compared to controls (Suppl. Fig. 4E-G). To prevent interference of aspecific antibodies in the assay, we compared the reactivity of IgM antibodies towards TOP1 and TOP1cc in the two ATA-IgM^+^ SSc patients and selected 9 other samples of ATA^+^ SSc patients with high levels of ATA-IgM from a previous study (Suppl. Fig. 4H, Table 1)^8^. TOP1cc was better recognized than TOP1 by IgM antibodies in these samples (Fig. 4E). The relative difference in the half-maximal binding titers to TOP1 and TOP1cc was comparable between IgG and IgM (Fig. 4F). Together, these data indicate that the differential recognition of TOP1 and TOP1cc is predominantly found in the IgM compartment and in the IgG1 subclass.

### ATA mAb can inhibit the enzymatic function of TOP1

We further characterized the ATA mAbs by assessing their effect on the enzymatic function of TOP1. Previous work showed that purified immunoglobulins from ATA^+^ SSc patients can inhibit the enzymatic function of TOP1^25,26^. The enzymatic activity of TOP1 can be measured based on the relaxation of supercoiled plasmid DNA which impairs the mobility of DNA through agarose gels (Fig. 5). Pre-incubation of TOP1 with ATA-IgG 2F8, ATA-IgG 7G6 and a control mAb did not affect the enzymatic function of TOP1 (Fig. 5). In contrast, supercoiled plasmid DNA was still apparent in the condition in which TOP1 was pre-incubated with ATA mAb 9D11 (Fig. 5). This indicates that ATA mAb 9D11 can inhibit the enzymatic function of TOP1.

**Figure 5.**
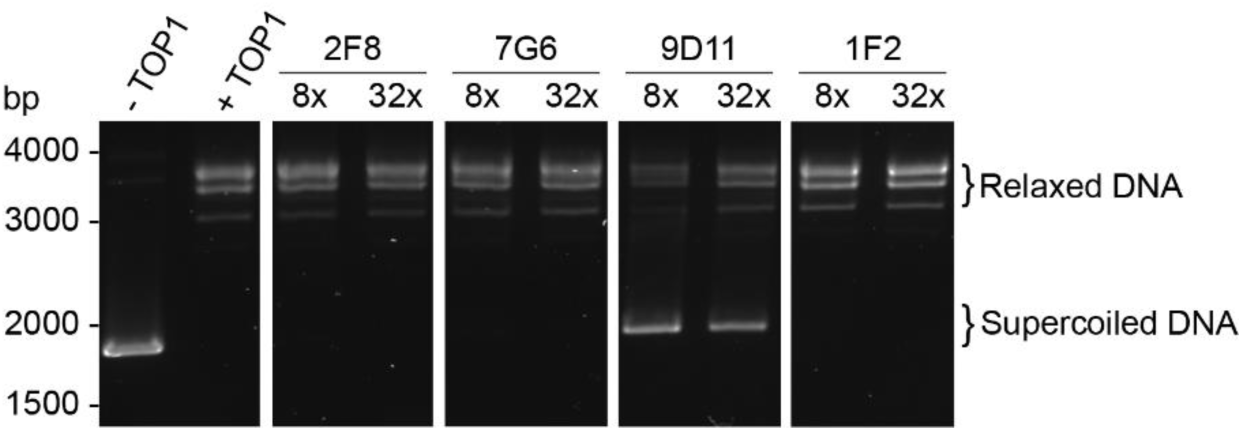
Effect of ATA mAbs on the enzymatic function of TOP1. DNA relaxation assay with TOP1 upon pre-incubation with an 8x and 32x molar equivalent of the ATA mAbs. A monoclonal antibody reactive towards citrullinated proteins (1F2) was used as negative control. DNA was visualized with a fluorescent DNA stain. Molecular weight of DNA in base pairs (bp) was estimated using a DNA ladder. Data are representative of two independent experiments.

### Enhanced induction of cytokine production by anti-TOP1cc mAbs in complex with TOP1cc

The enhanced binding of ATA mAbs to TOP1cc compared to TOP1 may have functional implications on the immunostimulatory capacity of ATAs. Therefore, we stimulated the monocytic THP-1 cell line with ATA mAbs complexed with plate-bound TOP1 and TOP1cc and subsequently determined the secretion of IL-8. Minimal secretion of IL-8 by THP-1 cells was detected in conditions with TOP1 and TOP1cc coating to which no antibody or a negative control antibody was added (Fig. 6, Suppl. Fig. 5). The ATA mAb 2F8, which binds to TOP1 and TOP1cc to a similar extent, induced comparable levels of IL-8 if complexed with TOP1cc or TOP1 (Fig. 6, Suppl. Fig. 5). In contrast, ATA-mAbs 7G6 and 9D11, which both bind TOP1cc better than TOP1, induced higher production of IL-8 by THP-1 when complexed with TOP1cc compared to TOP1 (Fig. 6, Suppl. Fig. 5). These data show functionality of the ATA mAbs generated and indicate that the ability of ATA to induce inflammation can be affected by the binding of TOP1 to DNA.

**Figure 6.**
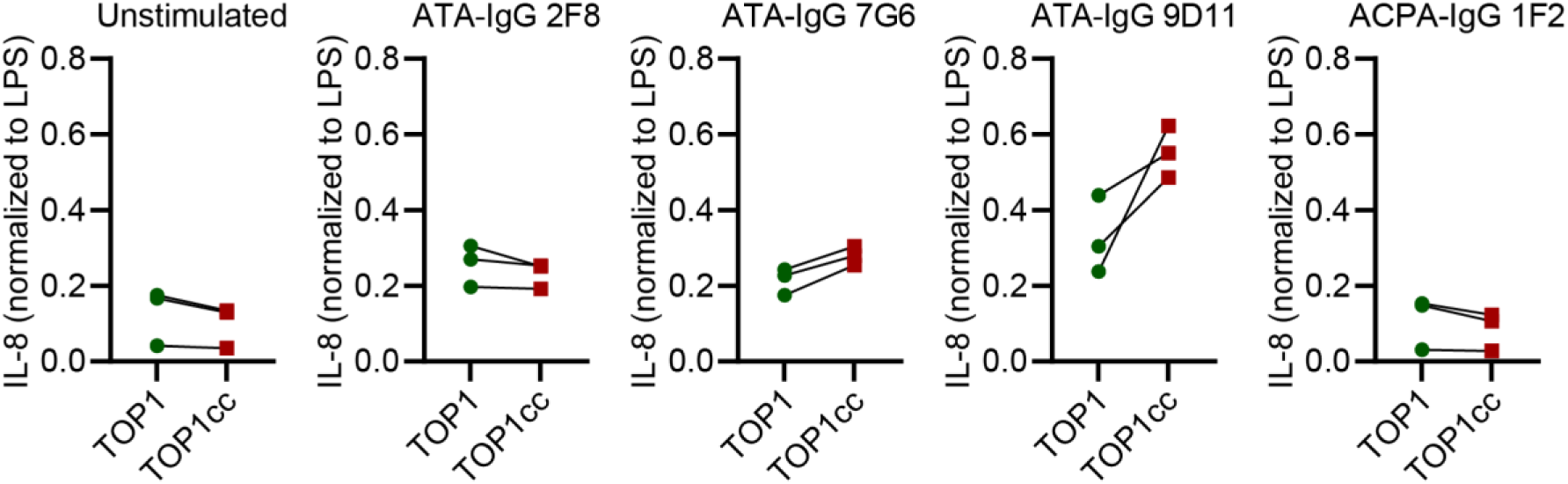
IL-8 production by THP-1 cells stimulated with ATA mAbs in complex with TOP1 and TOP1cc. Normalized levels of IL-8 in culture supernatants of THP-1 cells stimulated for 24 hours with ATA mAbs in complex with plate-bound TOP1 and TOP1cc. Conditions to which no mAb or the ACPA-IgG mAb 1F2 was added were used as negative controls. IL-8 levels were normalized to the levels of IL-8 detected in culture supernatant of THP-1 cells stimulated with 1µg/ml lipopolysaccharide (LPS). Data is obtained from three independent experiments which are connected by lines.

## Discussion

Predictability of disease progression forms a major challenge in the clinical management of SSc due to heterogeneity in clinical presentation and course of disease. To gain insights into the molecular mechanisms driving the disease, we here focused on the ATA B cell response and assessed whether DNA binding by TOP1 would affect its recognition by ATAs. By studying monoclonal and polyclonal ATAs, we observed remarkable heterogeneity among these autoantibodies in the recognition of TOP1 and TOP1cc. In fact, a subset of ATAs recognized TOP1cc better than TOP1, indicating that TOP1cc formation induces conformational changes, presumably, that enhance the antigenic properties of individual TOP1 epitopes and, consequently, the immunostimulatory properties of ATAs.

The differential recognition of TOP1 and TOP1cc by ATAs could relate to disease by various means. We observed that enhanced recognition of TOP1cc can be found in all ATA^+^ SSc patients, yet increasingly in patients with interstitial lung disease and in those with higher anti-TOP1-IgG levels in serum. Hence, it is likely that TOP1cc functions as an enhancer of the stimulation of ATA^+^ B cells. In fact, our data indicate that TOP1cc could preferentially stimulate a subset of B cells expressing ATA with enhanced recognition of TOP1cc epitopes. Stimulation of these cells may be further amplified by BCR-mediated internalization of DNA in complex with TOP1 and subsequent co-engagement of Toll-like receptor 9 (TLR9)^27–29^. Consequently, TOP1cc recognition may be relevant for both the initial break of B cell tolerance to TOP1 and, likewise, the continuous activation of ATA B cells in established disease. In this context, it is important that we noted the predominant presence of ‘TOP1cc-enhanced’ ATA within the IgM and the IgG1 subclass compartment. The presence of TOP1cc-enhanced reactivity in the IgM compartment makes the naive B cell compartment susceptible to strong BCR-mediated stimulation, while TLR stimulation synergizes with BCR signaling to induce AID which mediates BCR class-switching^28,30^. Previously, we could show that both the presence of ATA-IgM as well as ATA-IgG levels in serum associated with progressive disease, supporting this concept^8^. In addition, the presence of ATA-secreting cells in the circulation, a sign of recent B cell activation, associated with presence and severity of interstitial lung disease^7^. Together, these results indicate that the formation of TOP1cc could be important for ATA B cell activation and suggest that this process may be relevant to initiate and drive ATA^+^ disease.

Our findings imply that TOP1 in complex with DNA is the autoantigen primarily targeted by the ATA B cell response. Differential recognition of TOP1 and TOP1cc likely depends on conformational changes that occur when TOP1 binds DNA^31–33^. This conformational change only affects the binding of ATAs that recognize epitopes of TOP1 which are altered and, thus, probably represent a subset of ATA B cells. This is consistent with the finding that ATA mAbs differentially bind to TOP1 and TOP1cc, bind to unique epitopes and vary in their ability to inhibit the enzymatic function of TOP1. Also, our data indicate that such interaction may enhance ATA functional properties inducing or supporting inflammatory processes. The relevance of these functional effects and the context in which the actual antigen is encountered by ATAs and TOP1-reactive B cells remains to be determined, particularly within affected tissues. It has been described that TOP1 isolated from nuclei of SSc skin fibroblasts shows reduced enzymatic activity and is more often modified with a small ubiquitin-like modifier (SUMO)^34^. These traits are associated with the presence of TOP1cc, as TOP1cc is SUMOylated in pathways needed to repair DNA lesions trapped in TOP1cc^35^. Another variable could be the source of DNA incorporated in TOP1cc, as human TOP1 can also bind microbial DNA^36^. Microbial DNA is a more potent inducer of TLR9 signaling in comparison to human DNA, which might further enhance the pro-inflammatory potential of TOP1cc^37^. Together, such variables could determine the immunological properties of TOP1, add to the enhanced stimulation of B cells and indicate the need for molecular analyses of TOP1 in affected tissues.

In conclusion, our data indicate that DNA binding by TOP1, a nuclear autoantigen in SSc, affects the recognition of the antigen by disease-specific autoantibodies. This enhanced antigen binding extends the relevance of the well-appreciated pro-inflammatory effects of nucleic acid binding by nuclear antigens. Modulation of autoantigen recognition by nucleic acid binding could affect autoreactive B cell responses targeting nuclear proteins in SSc and other autoimmune diseases, thereby modulating their disease processes and course.

## Acknowledgements

Flow cytometry was performed at the Flow cytometry Core Facility (FCF) of the Leiden University Medical Center.

## Author contributions

Conceptualization: SN, DMHR, AJRH, CMF, JKVB, REMT, HUS

Designing and conducting experiments: SN, CW, DMHR, MJALW

Methodology: SN, CW, DMHR, MJALW, RQK, NL, CMF, REMT, HUS

Formal analysis: SN, DMHR, MJALW, HUS

Data interpretation: SN, DMHR, MJALW, AJRH, CMF, JKVB, REMT, HUS

Supervision: REMT, HUS

Writing original draft: SN, HUS

Revision of the manuscript and approval of the final version for submission: all authors.

