## Supplementary Figures for "Enhanced recognition of topoisomerase 1 upon DNA binding by a subset of anti-topoisomerase 1 autoantibodies in systemic sclerosis"

**Supplementary Figure 1.**

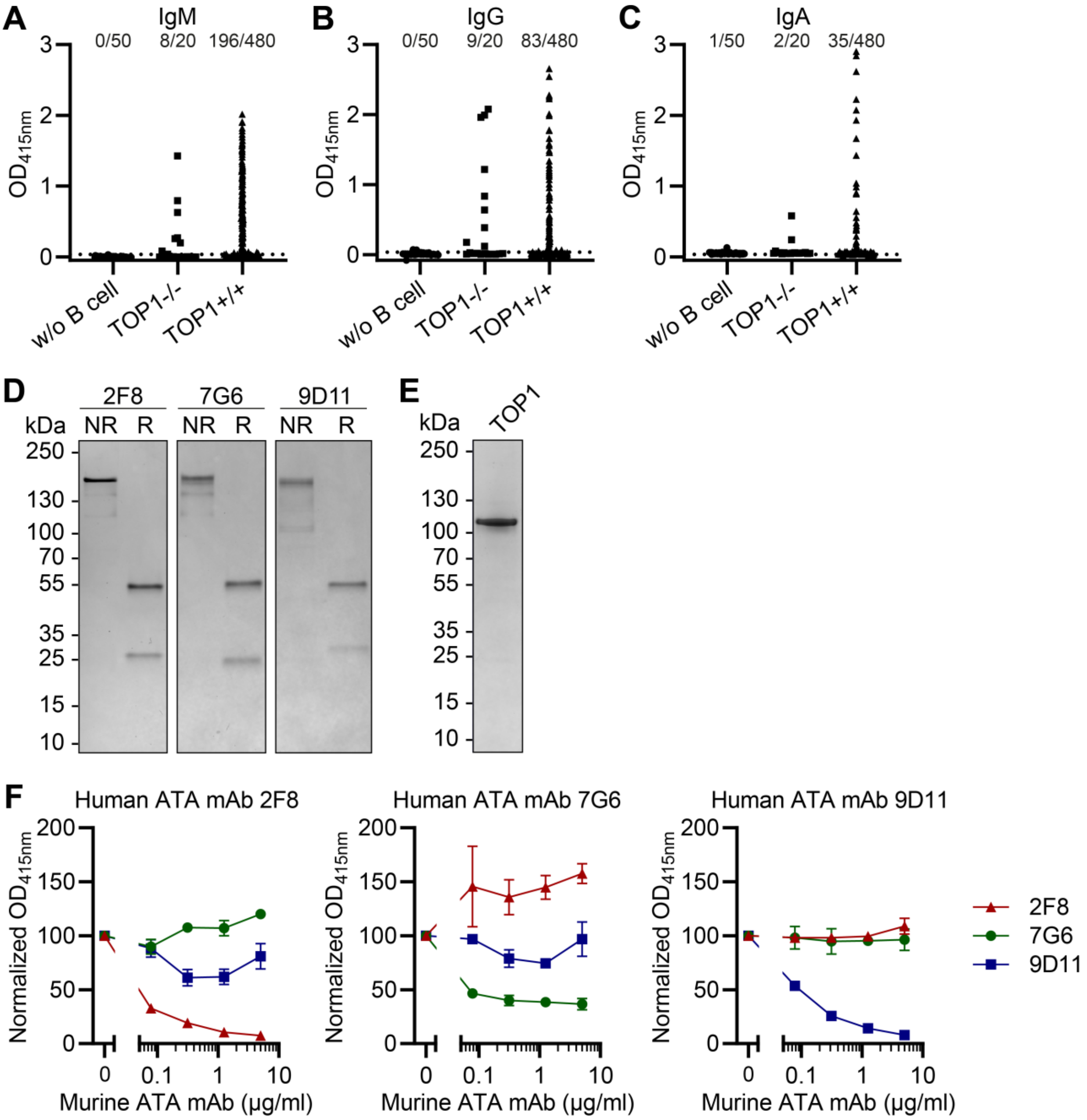


**Supplementary Figure 1. Generation and characterization of patient-derived ATA mAbs.**
**(A-C)** Presence of IgM (A), IgG (B) and IgA (C) in the culture supernatant of 480 single-cell sorted TOP1$^{+/+}$ B cells. Culture supernatants of 20 B cells lacking reactivity towards TOP1 (TOP1$^{-/-}$) and 50 culture supernatants without B cells (w/o B cells) were taken along as controls. Cut-off (mean of TOP1$^{-/-}$ B cells + 4x standard devation) is represented by the dotted line. Numbers indicate the amount of positive wells. **(D-E)** Gel electrophoresis of ATA mAbs under non-reducing (NR) and reducing (R) conditions (D) and of TOP1 under non-reducing conditions (E). Molecular weight in kilodaltons (kDa) was estimated using a protein ladder. **(F)** Binding of patient-derived ATA mAbs 2F8 (I), 7G6 (J) and 9D11 (K) with a human constant domain (human ATA mAbs) to TOP1-coated ELISA plates preincubated with ATA mAbs with a murine constant domain (murine ATA mAbs). Binding was normalized to the signal obtained when no murine mAb was added. Concentrations of human mAbs were in the linear range of the ATA-IgG ELISA; 2F8 - 0.04µg/ml, 7G6 & 9D11 - 1µg/ml. Data is representative of three independent experiments and each data point represents the mean and standard devation of two technical replicates from an individual experiment. (A-C, F) Optical density was measured at 415 nanometers ($OD_{415nm}$).

**Supplementary Figure 2.**

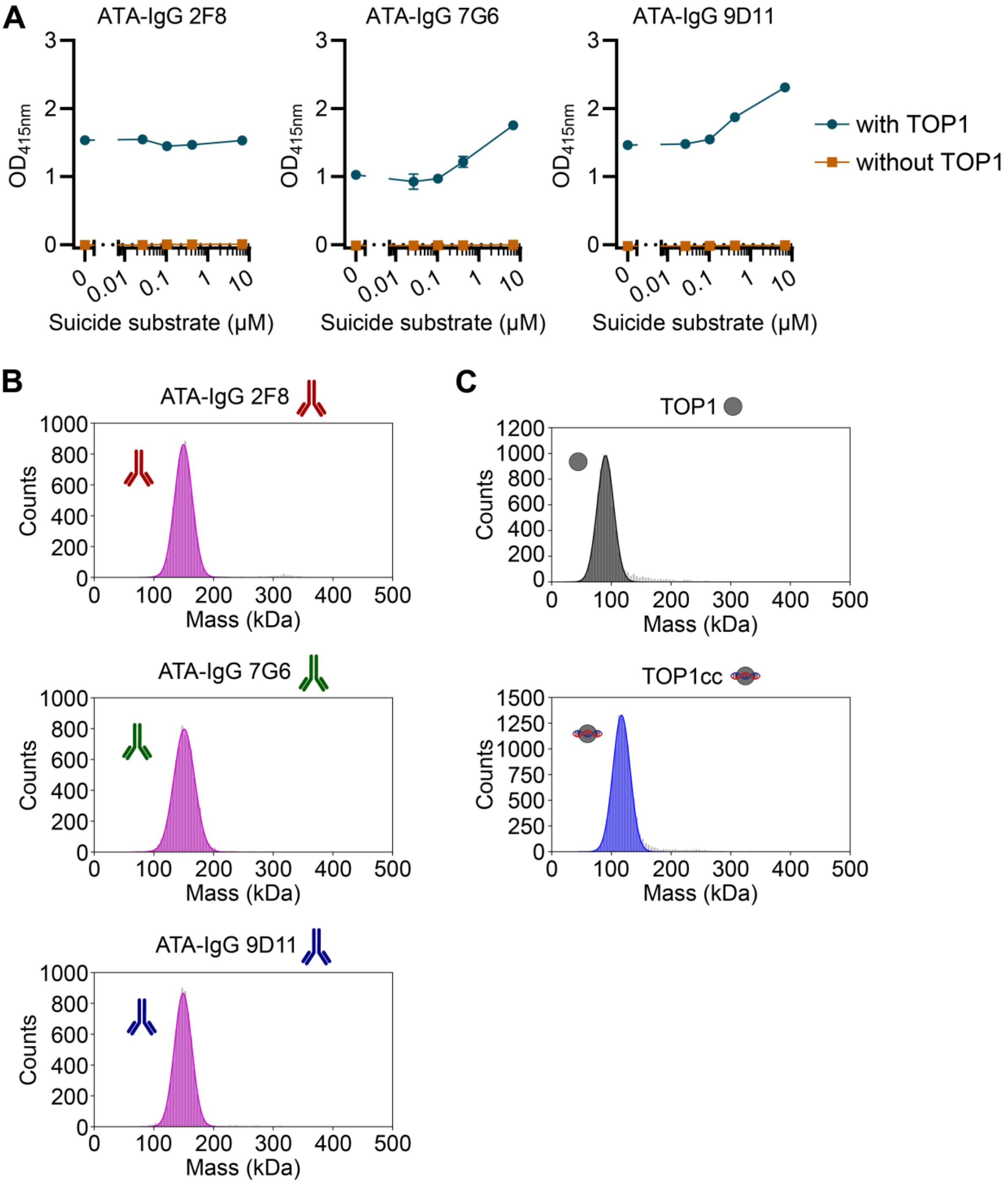


**Supplementary Figure 2. Validation of assays used to measure the reactivity of ATA mAbs towards TOP1 and TOP1cc by ELISA and mass photometry.**

**(A)** Binding of the ATA mAbs in an ELISA coated with and without TOP1 upon preincubation with a concentration range of suicide substrate. Concentrations of human mAbs were in the linear range of the ELISA; 0.08µg/ml of ATA-IgG 2F8, 9µg/ml of ATA-IgG 7G6 and 2.5µg/ml of ATA-IgG 9D11. Optical density was measured at 415 nanometers ($OD_{415nm}$). **(B-C)** Representative mass histograms of the ATA mAbs (B), TOP1 and TOP1cc (C) as obtained by mass photometry. An overview of detected masses can be found in Suppl. Table 1.

**Supplementary Figure 3.**

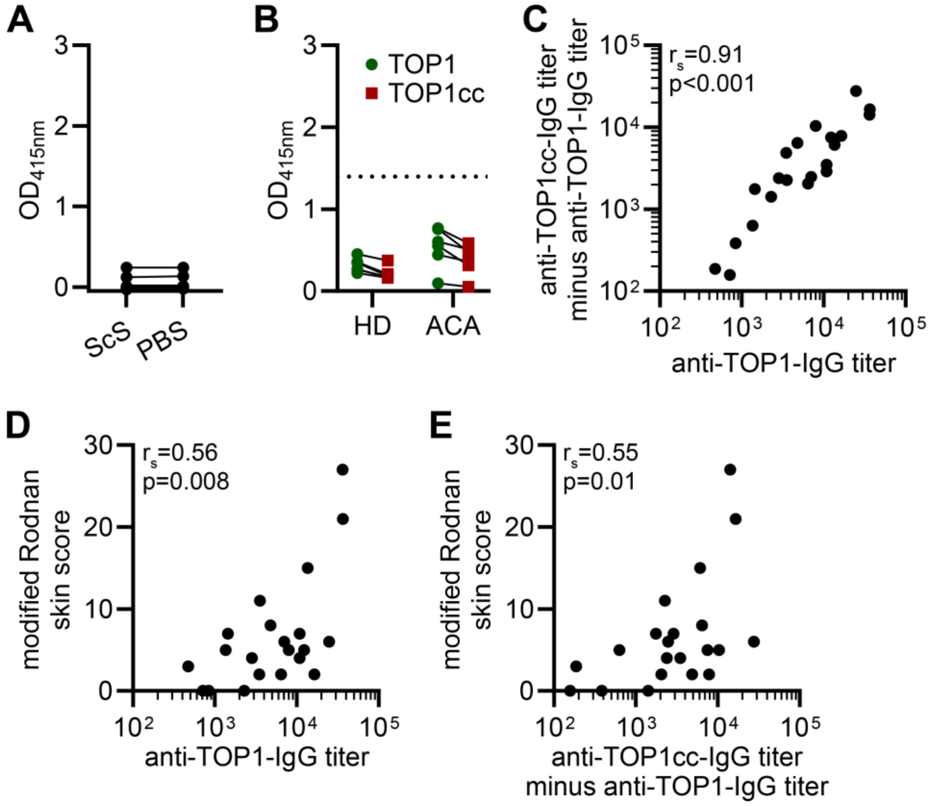


**Supplementary Figure 3. Relation between the recognition of TOP1 and TOP1cc and clinical disease.**
**(A)** Reactivity of IgG in plasma of ATA⁺ SSc patients (n=21) towards suicide substrate (ScS) and PBS (negative control). Signal was determined at the plasma concentration generating the half-maximal binding titer to TOP1 of the corresponding patient. **(B)** Reactivity of IgG in plasma of healthy donors (HD) (n=6) and ACA⁺ SSc patients (n=6) towards TOP1 and TOP1cc. Samples were diluted 625x, matching the lowest anti-TOP1-IgG half-maximal binding titers in a cohort of 21 ATA-IgG⁺ SSc patients. Half-maximal binding titer is represented by the dotted line (OD=1.4). **(C)** Correlation between anti-TOP1-IgG titer and difference between anti-TOP1cc-IgG and anti-TOP1-IgG titer (anti-TOP1cc-IgG titer minus anti-TOP1-IgG titer) in 21 ATA-IgG⁺ SSc patients. **(D-E)** Correlation between the modified Rodnan skin score (MRSS) and the anti-TOP1-IgG titer (H) and the difference between anti-TOP1cc-IgG and anti-TOP1-IgG titer. (A-B) Binding was based on optical density measured at 415 nanometers ($OD_{415nm}$). (C-E) Correlations are described using the Spearman's rank correlation coefficient.

**Supplementary Figure 4.**

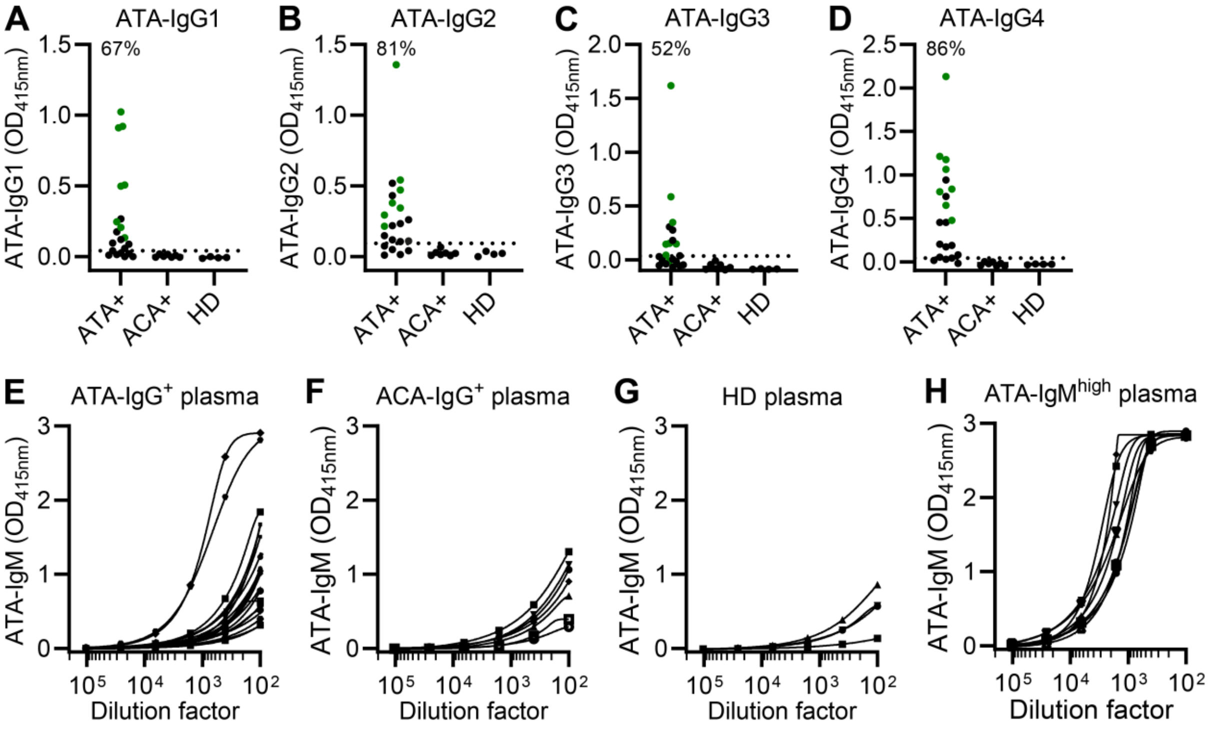


**Supplementary Figure 4. Binding of TOP1 by IgG subclasses and IgM in plasma of ATA$^+$ SSc patients.**

**(A-D)** Reactivity of IgG1-4 in plasma of ATA$^+$ SSc patients (n=21), ACA$^+$ SSc patients (n=7) and healthy donors (HD) (n=4) towards TOP1. Plasma samples were diluted 8000x for IgG1, 1000x for IgG2, 10000x for IgG3 and 3200x for IgG4. Cut-off (mean of ACA$^+$ SSc patients and HD's + 4x standard deviation) is represented by the dotted line. Frequency of positive samples are indicated in percentages. Selected samples used to analyze the reactivity of IgG subclasses towards TOP1 and TOP1cc are indicated in green. (**E-H**) Binding of TOP1 by IgM in serial diluted plasma of ATA$^+$ SSc patients (n=21) (E), ACA-IgG$^+$ patients (n=7) (F), healthy donors (HD) (n=4) (G) and ATA-IgM$^{high}$ patients (H). (A-H) Optical density was measured at 415 nanometers ($OD_{415nm}$).

**Supplementary Figure 5.**

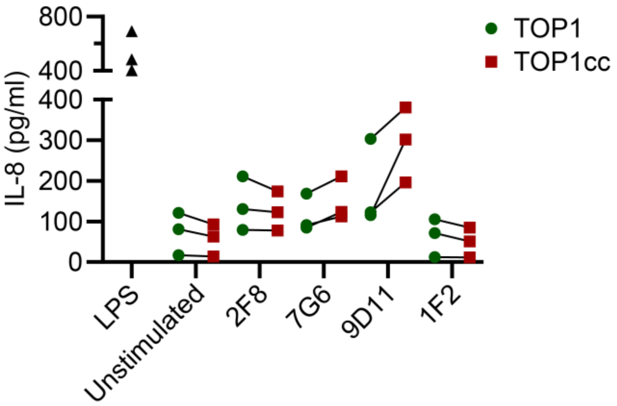


**Supplementary Figure 5. IL-8 concentration in culture supernatant of THP-1 cells stimulated with ATA mAbs in complex with plate-bound TOP1 and TOP1cc.**

IL-8 levels in culture supernatants of THP-1 cells stimulated for 24 hours with ATA mAbs in complex with plate-bound TOP1 and TOP1cc. Culture supernatants of THP-1 cells stimulated with 1μg/ml LPS were used as positive control. Conditions to which no mAb or ACPA-IgG mAb 1F2 was added were used as negative controls. Data is obtained from three independent experiments which are connected by lines.
