## Supplementary Methods for "Enhanced recognition of topoisomerase 1 upon DNA binding by a subset of anti-topoisomerase 1 autoantibodies in systemic sclerosis"

*Recombinant production and purification of TOP1*

Human TOP1 was recombinantly produced in insect cells (Protein Facility, Leiden University Medical Center) using an adapted Bac-to-Bac protocol (Invitrogen). In short, a pFastBacHT-A plasmid (OHu27246C, Genscript) encoding the full-length human TOP1 sequence with a N-terminal polyhistidine tag (6x histidine) and tobacco etch virus (TEV) protease recognition site (ENLYFQG) was commercially obtained and transformed into EmBACy competent cells (Geneva Biotech) to generate a recombinant bacmid. The recombinant bacmid was extracted from EmBACy cells by isopropanol precipitation and was subsequently used to transfect sedentary Sf9 insect cells (ThermoFisher). Upon culturing the transfected Sf9 cells for 72 hours at 28°C, culture supernatant containing baculovirus (P0) was collected. The baculovirus-containing supernatant was used to transduce suspension Sf9 cells to recombinantly express human TOP1 and create a baculovirus stock (P1). After culturing the cells for 72 hours at 28°C, transduced Sf9 cells were harvested and resuspended in His-buffer (50 mM HEPES pH7.5, 500 mM NaCl, 1% glycerol and 20 mM imidazole). Cells were lysed using sonication and TOP1 was purified using a HisTrap HP column (Cytiva), using a linear gradient of His-buffer supplemented with 500mM imidazole. Protein fractions were pooled and treated with TEV protease (Protein Facility, Leiden University Medical Center) under overnight dialysis against 20mM HEPES (pH=7.5), 200mM NaCl, 1mM EDTA and 1mM DTT. TOP1 was further purified by cation exchange chromatography (HiTrap SP FF, Cytiva) using a gradient between buffer A (20mM HEPES (pH=7.5), 50mM NaCl, 1mM EDTA and 1mM DTT) and buffer B (buffer A supplemented with 1M NaCl). Purity, integrity and antigenicity of recombinantly-produced TOP1 was validated by ELISA and gel electrophoresis using commercial TOP1 (ProSpec) as comparator.

*Generation of patient-derived anti-topoisomerase 1 monoclonal antibodies (ATA mAbs)*

ATA mAbs were generated based on the B cell receptor (BCR) sequence of TOP1-reactive B cells obtained from an ATA$^+$ SSc patient. To this end, peripheral blood mononuclear cells (PBMCs) were isolated by Ficoll-Paque gradient centrifugation. PBMCs were stained with LIVE/DEAD™ Fixable Aqua (Invitrogen), anti-CD3 BV510 (UCHT1, BioLegend), anti-CD14 BV510 (M5E2, BioLegend), anti-CD19 APC-Cy7 (SJ25C1, BD Biosciences), anti-CD20 FITC (2H7, BioLegend), anti-CD27 BV421 (M-T271, BD Biosciences) and TOP1 (Prospec) which was conjugated to PE (LNK021RPE, Biorad) and Alexa Fluor® 647 (A20006, Invitrogen) following the manufacturer's instructions. B cells binding to TOP1-PE and TOP1-Alexa Fluor® 647 were single-cell sorted with an FACS Aria III 4L (BD Biosciences) and subsequently cultured for 12 days as described previously[1]. Upon culturing, supernatants were harvested to analyze the antibodies secreted by the single-cell sorted B cells and cells were stored in Trizol (ThermoFisher Scientific) for BCR sequencing.

To determine whether secreted antibody was present in the culture supernatants of the single-cell cultured B cells, a total IgM, IgG and IgA ELISA was performed. High binding 384-well Microplates (Corning) were coated with 10µg/ml goat anti-human IgM, IgG or IgM (respectively A80-100, A80-104 and A80-102A, Bethyl Laboratories). Plates were blocked and supernatants were applied in a two-fold dilution. IgM, IgG and IgA were detected with 20ng/µl goat anti-human IgM-HRP, 50ng/ml goat anti-human IgG-HRP or 50ng/ml anti-human IgA-HRP (respectively A80-100P, A80-104P and A80-102P, Bethyl Laboratories). ABTS was used as substrate and plates were measured with a Spectramax I3x Multi-Mode Microplate Reader (Molecular Devices) using SoftMax® Pro 7 (version 7.0.2, Molecular Devices).

To determine the reactivity of the secreted antibodies to human TOP1, ATA-Ig, ATA-IgG and ATA-IgA ELISAs were performed. High binding 384-well Microplates (Corning) were coated with 2.5µg/ml human TOP1 in PBS (pH=8). Plates were blocked and culture supernatants were applied in a two-fold dilution. Ig, IgG and IgA was detected with goat anti-human Ig-HRP (A80-152P, Bethyl Laboratories), rabbit anti-human IgG-HRP (P0214, Dako) and goat anti-human IgA-HRP (A18781, Novex), respectively. ABTS was used as substrate and plates were measured with a Spectramax I3x Multi-Mode Microplate Reader (Molecular Devices) using SoftMax® Pro 7 (version 7.0.2, Molecular Devices).

The variable domain of the BCR of single cell sorted B cells which secreted human TOP1-reactive antibodies in culture were sequenced as described in detail previously[1]. In short, cells were lysed in Trizol and RNA was isolated. cDNA was generated from isolated RNA and used to perform the Anchoring Reverse Transcription of Immunoglobulin Sequences and Amplification by Nested (ARTISAN) PCR[2]. PCR products were sent for Sanger sequencing (Macrogen Europe) and BCR sequences were subsequently determined using IMGT-V-quest[3].

Based on the variable domain of the BCR sequences, plasmids were generated to recombinantly produce ATA mAbs. To this end, constructs containing the sequence encoding the variable domain of the ATA clones and a leader sequence were codon optimized (GeneArt, ThermoFisher Scientific) and ordered with the Kozak sequence (IDT DNA). These constructs were cloned into a pcDNA3.1 (+) expression vector (InVitrogen) using the In-Fusion® Cloning Kit (Takara Bio). To generate mAbs with human constant domains, constructs encoding heavy chain variable domains were cloned into a vector encoding the human IGHG1*03 constant region (P01857, UniProt) and constructs encoding light chain variable domains were cloned into a vector encoding the human IGLC3 or IGKC constant region (in accordance with the light chain of the original BCR; respectively P0DOY3 and P01834, UniProt). To generate mAbs with murine constant domains, constructs were cloned into vectors containing the murine IGHG2A*01 (V00825, IMGT Database) or murine IGLC1 or IGKC constant regions (respectively A0A0G2JE99 and P01837, UniProt).

The recombinant production of ATA mAbs by transfection of the vectors into FreestyleTM 293-F cells (Gibco) as well as the subsequent purification of IgG from culture supernatant using a 1ml HiTrap® Protein G HP affinity column (GE Healthcare) was performed as described previously[4]. A negative control human-derived citrullinated protein-reactive mAb (1F2) was obtained using a similar procedure as described above for the ATA mAbs[4]. Purity and integrity of the generated mAbs was validated by gel electrophoresis.

*Gel electrophoresis*

The purity and integrity of TOP1, TOP1cc and of the ATA mAbs was assessed by sodium dodecyl sulfate polyacrylamide gel electrophoresis (SDS-PAGE). TOP1 and TOP1cc were loaded on a polyacrylamide gel with a 4-15% gradient (Bio-Rad) upon addition of Laemmli buffer (Bio-Rad). The ATA mAbs were loaded on a polyacrylamide gel with a 4-15% gradient upon incubation for five minutes at 95°C in Laemmli buffer in presence (reduced) or absence (non-reduced) of 2% β-mercaptoethanol (Merck). PageRuler™ Plus Prestained Protein ladder (ThermoFisher Scientific) was used as reference. Proteins were separated for 60-90 minutes at 100-120 volt. Gels were subsequently washed in Milli-Q water for 5 minutes and stained for one hour using InstantBlue® Coomassie Protein Stain (Abcam). Upon overnight destaining in Milli-Q, gels were visualized using a ChemiDocTM Touch Imaging system (Bio-Rad).

*Western blotting*

Western blotting was used to determine the reactivity of ATA mAbs towards TOP1. Gel electrophoresis was performed as described above with 1µg of TOP1. Upon running the gel, gels were blotted on a polyvinylidene difluoride membrane (Bio-Rad). Membrane was blocked with 3% skim milk powder (Sigma) in 0.05% Tween in PBS. Subsequently, 1µg/ml of ATA mAb was added to the membrane and incubated overnight at 4°C. Upon washing the membrane, rabbit anti-human IgG (1:2000, P0214, DAKO) was used as secondary antibody. Washed membranes were developed using Pierce™ enhanced chemiluminescence substrate (ThermoFisher Scientific) and visualized using a ChemiDoc™ Touch Imaging system (Bio-Rad).

*Cross-inhibition ELISA with ATA mAbs*

Cross-inhibition ELISAs using ATA mAbs with murine and human constant domains were performed to determine whether ATA mAbs can affect the binding of other ATA mAbs to TOP1. To this end, the anti-TOP1-IgG ELISA was adapted by the addition of ATA mAbs with murine constant domains in a serial dilution to the plates before sample addition. Upon washing the plates, ATA mAbs with human constant domains were added and the anti-TOP1-IgG ELISA was conducted as described above.

*Mass photometry*

Mass photometry was used to determine the stoichiometry of the interaction between the ATA mAbs to TOP1 and TOP1cc. ATA mAbs were incubated at a concentration of 100nM with an excess of TOP1 or TOP1cc for five minutes at room temperature. Subsequently, samples were diluted fivefold in PBS and analyzed using a OneMP mass photometer (Refeyn) as described previously[5,6]. The analysis was conducted on custom-prepared sample carrier slides made from reusable cell culture gaskets (Grace Biolabs) and microscope coverslips (24mm x 50mm, Paul Marienfeld GmbH). Samples were recorded with AcquireMP software (Refeyn) for 60 seconds at a scan rate of 90 fragments per second using the medium field of view. In addition to the protein mixtures, ATA mAbs, TOP1 and TOP1cc were also measured separately at a final concentration of 20nM to determine the molecular mass of the individual components. An in-house standard mix consisting of IgM (1006 kDa), apoferritin (513 kDa), IgG (149 kDa), IgG half-body (73 kDa) was used to perform mass calibration. Calibrated data were then exported and processed using a custom Python pipeline, incorporating seaborn, SciPy, NumPy, pandas and Matplotlib[7–11]. Mass histograms were generated with 5kDa bin widths and resulting mass distributions were manually annotated. SciPy was further used to analyze the mass histograms, to identify peaks corresponding to the different molecular species, and to estimate the molecular weights of the observed peaks.

*TOP1-mediated DNA relaxation assay*

A TOP1-mediated DNA relaxation assay was used to study the effect of the ATA mAbs on the enzymatic function of TOP1. To this end, a 10µl suspension containing 1µg/ml TOP1 with a 8x or 32x molar equivalent of ATA mAb in reaction buffer (40mM Tris-HCl, 300mM KCl, 2mM EDTA, 2mM DTT, 100µg/ml BSA) was incubated for one hour at room temperature. Subsequently, 10µl of 20µg/ml negatively supercoiled pUC19 DNA plasmid (ThermoFisher Scientific) was added before performing the enzymatic reaction for 30 minutes at 37°C. Afterwards, 10µl of 2.5% SDS was added and the mixture was incubated at 37°C for 15 minutes. Reaction mixture was mixed with Orange G loading dye (Merck) and ran on a 1% agarose gel at 100 volt for 2-3 hours. GeneRuler 1kb Plus DNA ladder (ThermoFisher Scientific) was used as reference. DNA in the gel was stained using Nancy-520 (Merck). Gel was visualized using a ChemiDoc™ MP Imaging system (Bio-Rad).
