## Supplementary Table for "Enhanced recognition of topoisomerase 1 upon DNA binding by a subset of anti-topoisomerase 1 autoantibodies in systemic sclerosis"

**Supplementary Table 1. Overview of experimental masses determined by mass photometry.**

| Sample | Species | Experimental mass (kDa) | Confidence interval (standard deviation) | Theoretical mass (kDa) |
|---|---|---|---|---|
| TOP1 | TOP1 | 90 | 13 | 91 |
| TOP1cc | TOP1cc | 116 | 14 | 119 |
| ATA-IgG 2F8 | mAb | 149 | 14 | 144 |
| ATA-IgG 7G6 | mAb | 150 | 17 | 145 |
| ATA-IgG 9D11 | mAb | 148 | 14 | 145 |
| ATA-IgG 2F8 + TOP1 | TOP1 | 93 | 12 | 91 |
| | mAb + 2x TOP1 | 352 | 17 | 326 |
| ATA-IgG 2F8 + TOP1cc | TOP1cc | 114 | 14 | 119 |
| | mAb + 2x TOP1cc | 392 | 21 | 382 |
| ATA-IgG 7G6 + TOP1 | TOP1 | 91 | 13 | 91 |
| | mAb + 1x TOP1 | 247 | 22 | 236 |
| | mAb + 2x TOP1 | 343 | 18 | 327 |
| ATA-IgG 7G6 + TOP1cc | TOP1cc | 115 | 13 | 119 |
| | mAb + 2x TOP1cc | 399 | 19 | 383 |
| ATA-IgG 9D11 + TOP1 | TOP1 | 89 | 11 | 91 |
| | mAb | 140 | 15 | 145 |
| | mAb + 1x TOP1 | 237 | 17 | 236 |
| | mAb + 2x TOP1 | 337 | 21 | 327 |
| ATA-IgG 9D11 + TOP1cc | TOP1cc | 114 | 15 | 119 |
| | mAb + 1x TOP1cc | 250 | 23 | 264 |
| | mAb + 2x TOP1cc | 387 | 21 | 383 |
